# Combining In Vivo Corneal Confocal Microscopy with Deep Learning-based Analysis Reveals Sensory Nerve Fiber Loss in Acute SIV Infection

**DOI:** 10.1101/2020.04.19.048926

**Authors:** Megan E. McCarron, Rachel L. Weinberg, Jessica M. Izzi, Suzanne E. Queen, Stuti L. Misra, Daniel B. Russakoff, Jonathan D. Oakley, Joseph L. Mankowski

**Author notes:** Correspondence: Joseph L Mankowski, Department of Molecular and Comparative Pathobiology, Johns Hopkins University, Baltimore, MD. Contributed equally (Megan E. McCarron, Rachel L. Weinberg).

## Abstract

**Purpose:** To characterize corneal subbasal nerve plexus morphologic features using *in vivo* corneal confocal microscopy (IVCM) in normal and SIV-infected macaques and to implement automated assessments using novel deep learning-based methods customized for macaque studies.

**Methods:** In vivo corneal confocal microscopy images were collected from both male and female age-matched specific-pathogen free rhesus and pigtailed macaques housed at the Johns Hopkins University breeding colony using the Heidelberg HRTIII with Rostock Corneal Module. We also obtained repeat IVCM images of 12 SIV-infected animals including pre-infection and 10 day post-SIV infection time-points. All IVCM images were analyzed using a novel deep convolutional neural network architecture developed specifically for macaque studies.

**Results:** Deep learning-based segmentation of subbasal nerves in IVCM images from macaques demonstrated that corneal nerve fiber length (CNFL) and fractal dimension measurements did not differ between species, but pigtailed macaques had significantly higher baseline corneal nerve fiber tortuosity than rhesus macaques (P = 0.005). Neither sex nor age of macaques was associated with differences in any of the assessed corneal subbasal nerve parameters. In the SIV/macaque model of HIV, acute SIV infection induced significant decreases in both corneal nerve fiber length and fractal dimension (P= 0.01 and P= 0.008 respectively).

**Conclusions:** The combination of IVCM and objective, robust, and rapid deep-learning analysis serves as a powerful noninvasive research and clinical tool to track sensory nerve damage, enabling early detection of neuropathy. Adapting the deep-learning analyses to human corneal nerve assessments will refine our ability to predict and monitor damage to small sensory nerve fibers in a number of clinical settings including HIV, multiple sclerosis, Parkinson’s disease, diabetes, and chemotherapeutic neurotoxicity.

## Introduction

Human immunodeficiency virus (HIV) associated sensory neuropathy remains a common and debilitating complication of HIV infection, even with antiretroviral therapy (ART).(1, 2) Signs of HIV neuropathy can develop early during primary HIV infection before most patients start antiretroviral therapy.(3) Studies in SIV/macaque models of HIV PNS damage have demonstrated that SIV infects and replicates within macrophages in sensory ganglia during the first week of infection followed shortly thereafter by immune activation of macrophages, at day 10 post-infection.(3-6) Concordant loss of epidermal nerve fibers in the skin of the hind feet can be detected at 14 days post-SIV infection.(7)

The current gold standard to measure sensory nerve fiber loss in both research and clinical settings is to calculate epidermal nerve fiber density in skin biopsies after PGP9.5 immunostaining to identify the small sensory nerve fibers.(8) We have adapted and validated similar approaches for use in macaques; however, serial biopsies are problematic for animals and human subjects, as it requires repeat sampling within the same local region of sensory innervation. Accordingly, we have developed an alternative non-invasive approach to measure sensory fiber loss amenable to longitudinal tracking of individual animals using *in vivo* corneal confocal microscopy (IVCM). This technique has previously been utilized in human studies as a tool to detect early changes in corneal subbasal nerves prior to diabetic peripheral neuropathy and has great potential for tracking sensory fiber loss longitudinally in HIV patients.(3, 9-13). Therefore, IVCM can be used as a marker to potentially diagnose and monitor progression of peripheral neuropathy. Our prior work using βIII tubulin immunostained corneal whole mount sections coupled with automated analysis first demonstrated that alterations in nerve fiber density of the subbasal plexus of the cornea correlated with epidermal nerve fiber loss in SIV animals that progressed to AIDS.(14) This work extends these methods and approach to *in vivo* imaging. The aims of the current study were to establish reference normal values for macaques and to investigate potential SIV-induced changes in the corneal subbasal nerve plexus using IVCM combined with novel deep learning-based methods customized for image processing for macaque studies.

## Materials and Methods

### Animals

All uninfected macaques used in this study were housed in indoor-outdoor enclosures in harem breeding groups at the Johns Hopkins University (JHU) Research Farm. Pigtailed macaques (*Macaca nemestrina*) and rhesus macaques (*Macaca mulatta*) were housed separately. Animals were screened annually and consistently tested serologically negative for *Macacine herpesvirus* 1 (B virus), simian immunodeficiency virus (SIV), simian T-cell leukemia virus (STLV-1), and simian retrovirus (SRV), comprising the battery of pathogens excluded to establish specific-pathogen free (SPF) status. Intradermal tuberculin skin tests were performed semi-annually and all animals in the colony were consistently negative.

### Animal study groups

IVCM images were obtained from 21 healthy rhesus macaques (10 females and 11 males) and 40 pigtailed macaques (17 females and 23 males) to establish normal reference values for corneal nerve fibers that form the subbasal plexus. To determine whether age was associated with alterations in corneal subbasal nerve fibers (SNF), macaques were separated into four age groups (3y, 4 to 6y, 7 to 10y, and over 10 y); each age group was composed of 10 to 23 animals. Animals included in the study had no history or current clinical evidence of ocular disease, neurologic disease, or systemic diseases known to cause neurologic sequelae, such as diabetes mellitus.

For SIV studies,12 juvenile pigtailed macaques co-housed at JHU were inoculated intravenously with SIV/17E-Fr and SIVdeltaB670.(15) Animals had corneal confocal scans performed under ketamine sedation 10 to14 days before SIV inoculation and then at 10 days post-inoculation to determine whether corneal sensory nerve fiber alterations could be detected during acute SIV infection. All animal work was approved by the JHU IACUC and determined to be in accordance with the guidelines outlined in the Animal Welfare Act and Regulations (United States Department of Agriculture) and the 8^th^ edition of the Guide for the Care and Use of Laboratory Animals (National Institutes of Health).

### Image acquisition

Central corneal images were acquired from all animals using the Heidelberg Retina Tomograph III (HRT III) Rostock Cornea Module (Heidelberg Engineering, Heidelberg, Germany). Each image covered a field of view of 400μm by 400μm over 384 by 384 pixels. Animals were anesthetized with 10mg/kg ketamine HCl (Zetamine, VetOne, Boise, ID) intramuscularly for imaging. If needed, animals received additional doses of either 5-10mg/kg ketamine HCl intramuscularly or intravenously, 0.1-0.5mg/kg acepromazine (Acepromazine Maleate Injection, USP, VetOne, Boise, ID) intramuscularly, or 0.1-0.2mg/kg midazolam (Midazolam Injection, USP, Hospira, San Jose, CA) intramuscularly to extend anesthesia time. Exact drug and dosage regimens were determined by the veterinarian for each animal to maintain sedation for the duration of the imaging session. Once anesthetized, 1-2 drops of 0.5% proparacaine hydrochloride ophthalmic solution (Alcon, Fort Worth, TX) were applied topically to the right eye of each animal. A thin layer of a sterile lubricant eye gel (GenTeal Lubricant Eye Gel, Alcon, Fort Worth, TX) was also applied to the right eye of each animal and a speculum was placed to retract the eyelids throughout the imaging session. A thin layer of sterile lubricant eye gel was applied to the tip of the objective lens and a sterile disposable cap (Tomocap, Heidelberg Engineering, Heidelberg, Germany) was placed over the lens. The head of the anesthetized animal was manually positioned to orient the eye towards the corneal surface. Images of the subbasal nerve plexus of the cornea were acquired from each animal using a sequence scan. Imaging was performed until 5 high-quality images of the subbasal nerve plexus (SBP) in the central cornea were obtained. Each imaging session lasted between 5 to 15 minutes.

### Measuring Plasma SIV RNA

SIV RNA was measured by qRT-PCR using the QuantiTech Virus Kid (Qiagen) and the following primers and probes in the SIV gag region (Fwd: 5’GTCTGCGTCATCTGGTG CATTC-3’, Rev: 5’-CACTAGGTGTCTCTGCAC TATCTGTTTTG-3’, Probe: 5’-CTTCCTCAGTGTGTTTCACTTTCTCTTCTG-3’) with the following cycling conditions 50°C for 30 min to reverse transcribe RNA, 94°C for 15 min, which was followed by 45 cycles of PCR at 94°C for 15 seconds, 55°C for 15 seconds, and 60°C for 30 seconds as previously described.(5)

### Deep Learning-Based Analysis: Applying deepNerve for automated assessments

As previously described in detail, we developed a customized deep learning-based approach termed deepNerve to measure objectively three parameters of interest in the macaque subbasal plexus: corneal nerve fiber length (CNFL), fractal dimension, and tortuosity. (16) This algorithm was specifically built using macaque corneal nerve images obtained by IVCM as current automated approaches designed to assess the SBP were developed for human studies. Image quality from macaque IVCM scans lags behind human image quality because 1) macaque eyes are smaller with a more acute radius of curvature and a thinner cornea leading to a lower signal to noise image in the periphery of the images, and 2) macaques need to be anesthetized and manually positioned for IVCM unlike compliant human subjects who can be directed to aid orientation of the cornea to the microscope objective. For each individual animal corneal scan, the 5 highest quality images of subbasal plexus were selected for deepNerve input. These highest quality corneal scan images from macaques were processed to obtain three distinct parameters:

1. Corneal nerve fiber length – the total lengths of the nerves divided by the area of the image (mm/mm^2^). (17)
2. Fractal dimension - a measure of nerve complexity that assesses the nerve pattern over scale and records the spatial loss of nerves. We used the box counting method over the same scales as previously published. (18)
3. Tortuosity - the Euclidian distance between the end points of each nerve segment divided by the true length of that segment. The more tortuous the vessel is over its extent, the longer the traversable length relative to the Euclidian distance and the higher the metric. Nerve branch points are also included as end points defining a nerve segment. (16)

## Data Analysis

The median values for all parameters calculated by deepNerve for each individual corneal confocal scan were used as representative values for subsequent data analysis for each animal. Group comparisons between species and sex were performed using the t-test; age comparisons utilized ANOVA. The changes from baseline pre-infection to 10 days post-SIV infection within individual animals were analyzed using the paired t-test. All statistical analyses were performed using GraphPad Prism v. 8.0 with α = 0.05.

## Results

While a recent report illustrated the salient architectural features of subbasal nerve fibers in macaques based on immunohistochemical staining of corneal whole mounts, corneal subbasal plexus nerve fiber morphology in IVCM images from macaques has not been reported previously.(19) In this study, visual comparison of corneal confocal scans obtained from a human subject (51 year old male), a pigtailed macaque (4 year old male), and a rhesus macaque (7 year old female) revealed similar nerve morphology in the subbasal plexus of the central cornea (Figure 1). All primate species exhibited comparable subbasal nerve plexus architecture with predominantly parallel arrangement of nerve fibers, and both branching of nerve fibers and interconnecting smaller diameter anastomoses between the main nerve fibers. Macaque nerves in general were observed to be thinner than human nerves.

**Figure 1.**
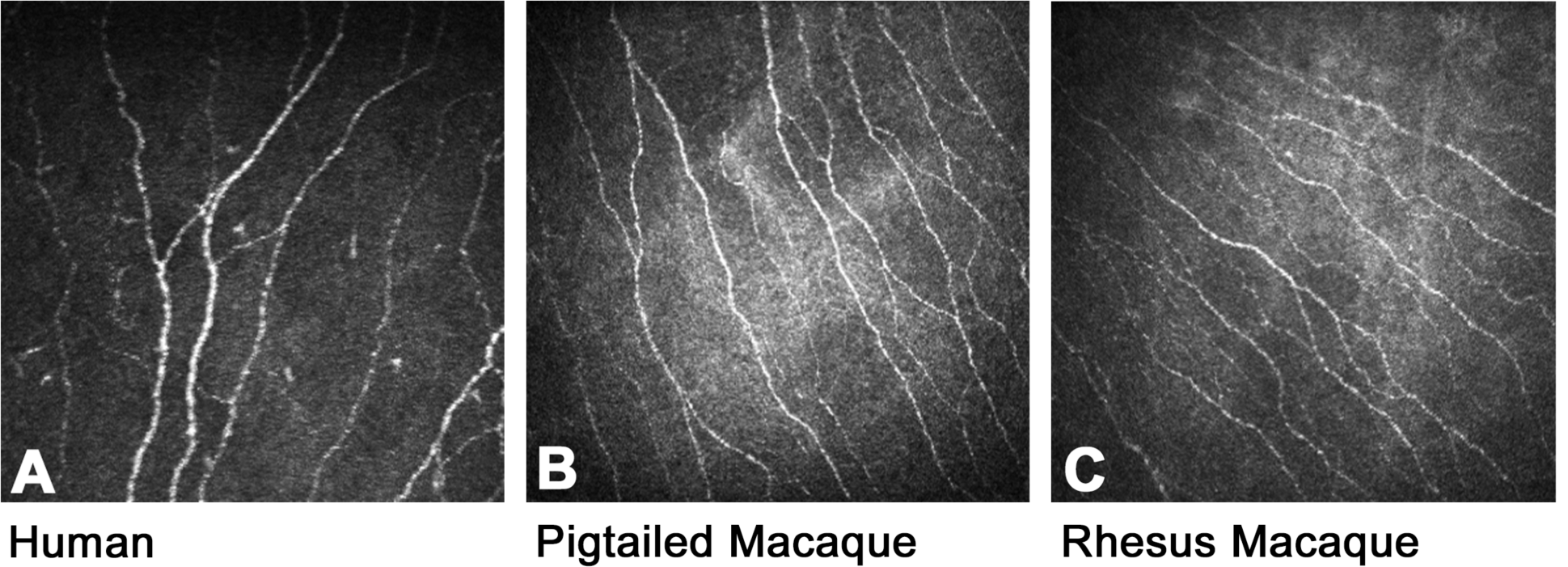
Representative images of the central corneal subbasal nerve plexus obtained by in vivo corneal confocal microscopy from A) a human, B) a pigtailed macaque, and C) a rhesus macaque illustrates similar density, arrangement, and morphology of the small sensory nerves across these primate species. Each image covers the same area of 400µm-by-400µm.

Corneal confocal scan images often have been assessed using manual tracing techniques that are inherently subjective and time-consuming. In contrast, nerve fiber detection by a customized deep learning-based approach rapidly and objectively identifies the large majority of individual nerve fibers in scans, facilitating subsequent automated measurements of nerve fiber parameters. This approach, termed deepNerve, was specifically designed to identify nerves present in macaque corneal scans as there is no current automated platform available for this purpose in macaques. (16)

After deepNerve processing, we compared corneal nerve parameters between healthy pigtailed macaques and rhesus macaques housed at JHU. There are notable differences between these species with regard to sensory nerve fiber innervation of the epidermis and SIV outcomes including extent of nervous system damage, thus establishing and comparing normative data is extremely valuable to enhance the rigor and reproducibility of macaque-based research.(20) There were no significant differences identified between normal pigtailed macaques versus comparable rhesus macaques for corneal nerve fiber length or fractal dimension; however, nerve fiber tortuosity was significantly higher in pigtailed macaques than rhesus macaques (P = 0.005; Fig 2; Table 1).

**Table 1.** Normative values for macaque IVCM corneal nerve parameters

|  | CNFL (mm/mm<sup>2</sup>) |  | Fractal Dimension |  | Tortuosity |  |
| --- | --- | --- | --- | --- | --- | --- |
| **Species** | MEAN | SD | MEAN | SD | MEAN | SD |
| Pigtailed | 20.10 | 4.46 | 1.27 | 0.05 | 1.025 | 0.005 |
| Rhesus | 19.96 | 4.68 | 1.27 | 0.05 | 1.021 | 0.005 |
| **Sex** |  |  |  |  |  |  |
| Female | 20.88 | 3.97 | 1.28 | 0.04 | 1.023 | 0.005 |
| Male | 19.35 | 4.84 | 1.26 | 0.05 | 1.023 | 0.005 |
| **Age** |  |  |  |  |  |  |
| 0-3 yrs | 18.87 | 4.15 | 1.26 | 0.05 | 1.022 | 0.004 |
| 4-6 yrs | 20.29 | 5.20 | 1.28 | 0.01 | 1.023 | 0.005 |
| 7-10 yrs | 21.68 | 4.51 | 1.29 | 0.01 | 1.022 | 0.006 |
| >10 yrs | 19.89 | 4.15 | 1.27 | 0.02 | 1.027 | 0.005 |

**Figure 2.**
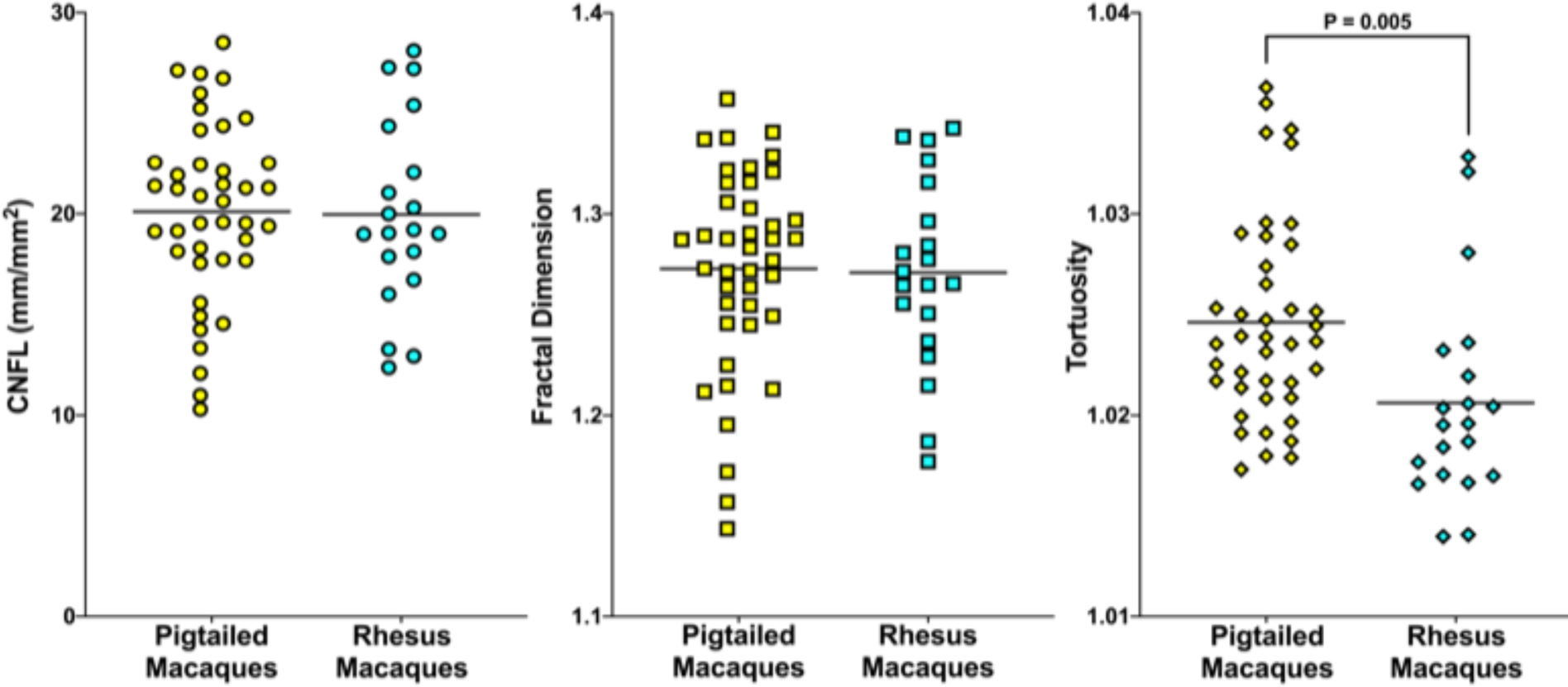
CNFL (left) and Fractal Dimension (center) did not differ significantly between pigtailed macaques versus rhesus macaques. However, a significantly higher tortuosity was observed in pigtailed macaques compared with rhesus macaques (right).

As sex can be a major variable for many biological measurements, comparisons of corneal nerve parameters were made between female macaques versus male macaques for all corneal measurements. No significant differences were identified in these comparisons, suggesting that baseline corneal nerve parameters do not differ between males and females (Figure 3, Table 1), similar to reports in humans.(21)

**Figure 3.**
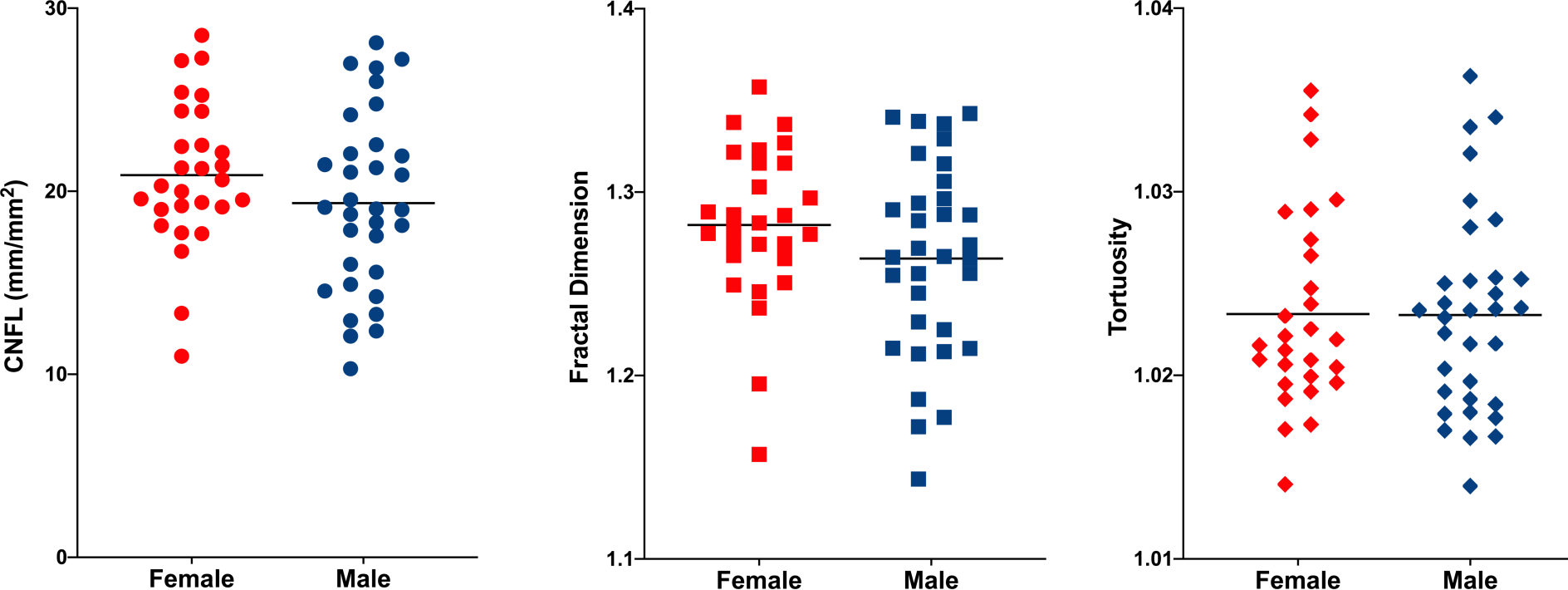
CNFL (left), Fractal Dimension (center), and Tortuosity (right) did not differ significantly between male versus female macaques.

The influence of age on corneal nerve parameters was examined after categorizing healthy animals into one of four age groups, (Figure 4, Table 1). Although there was a trend for tortuosity to increase with age in animals older than 10 years, this was not a significant finding. As we had fewer animals in the older age group, it is possible that a larger scale study might find a significant increase with maturation. Animals past breeding age were not included in this study as it was based in the JHU breeding colony; follow up studies in aging macaque cohorts may reveal age-related alterations in corneal sensory innervation. Of note, reduced corneal nerve fiber density has been reported with aging in both mice and humans of advanced age. (21, 22)

**Figure 4.**
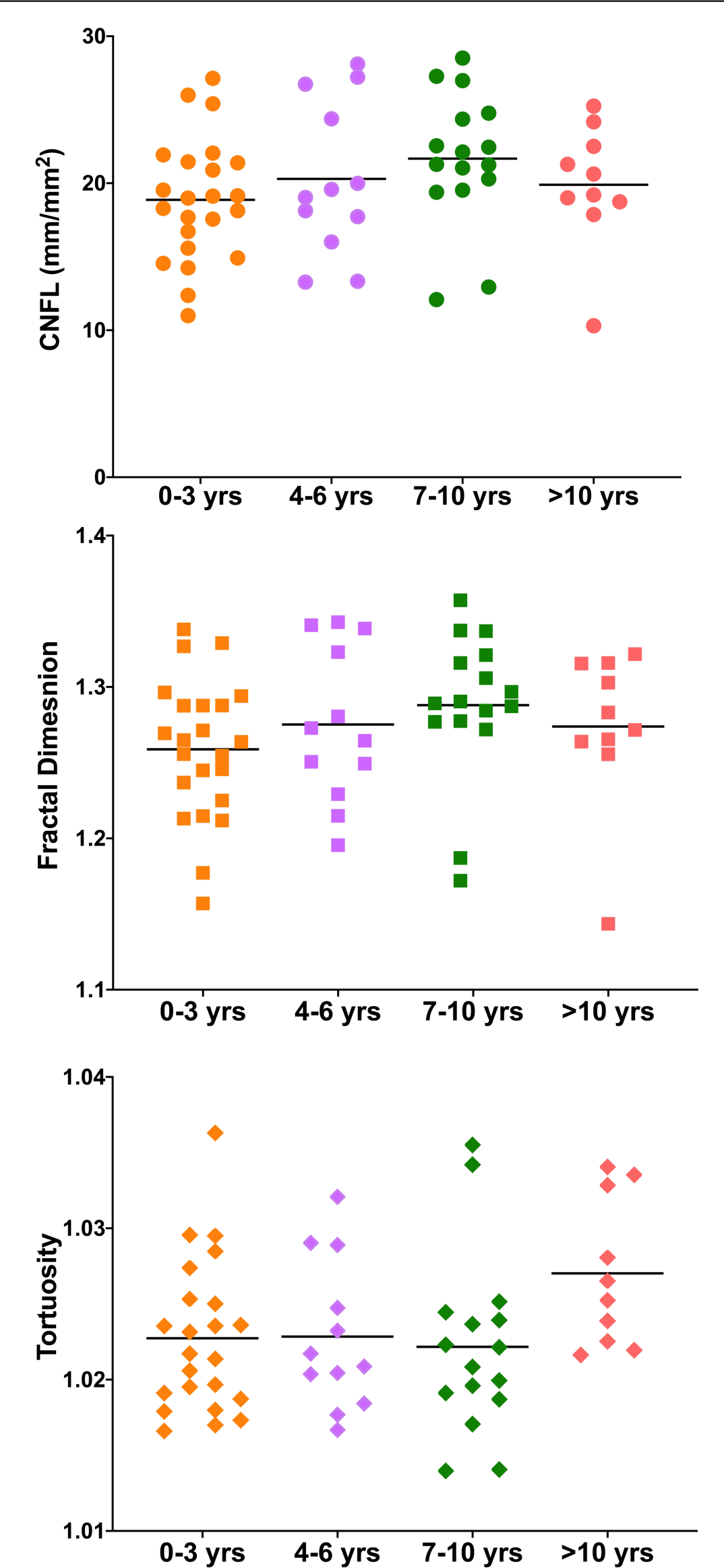
CNFL (top), Fractal Dimension (center), and Tortuosity (bottom) stratified by age. Only tortuosity was higher in the oldest group, but not significantly.

Our previous studies of HIV-induced damage to the peripheral nervous system based in the SIV/pigtailed macaque model have revealed significant loss of small sensory nerve fibers in the epidermis of the skin in the SIV/pigtailed macaque model at 14 days post-SIV infection (25); similar early loss of epidermal nerve fibers was found in a rhesus macaque SIV model 21 days post-SIV infection.(23) In this study, to determine whether IVCM-based analysis could detect early sensory nerve fiber damage, we performed IVCM on 12 pigtailed macaques to obtain pre-infection baseline corneal nerve data followed by IVCM scans on day 10 days after infection with SIV. All infected animals had similar plasma viral loads (Fig 5). A decline in CNFL density was visually apparent when pre-infection IVCM scans and deepNerve processed images were compared (Fig 6).

**Figure 5.**
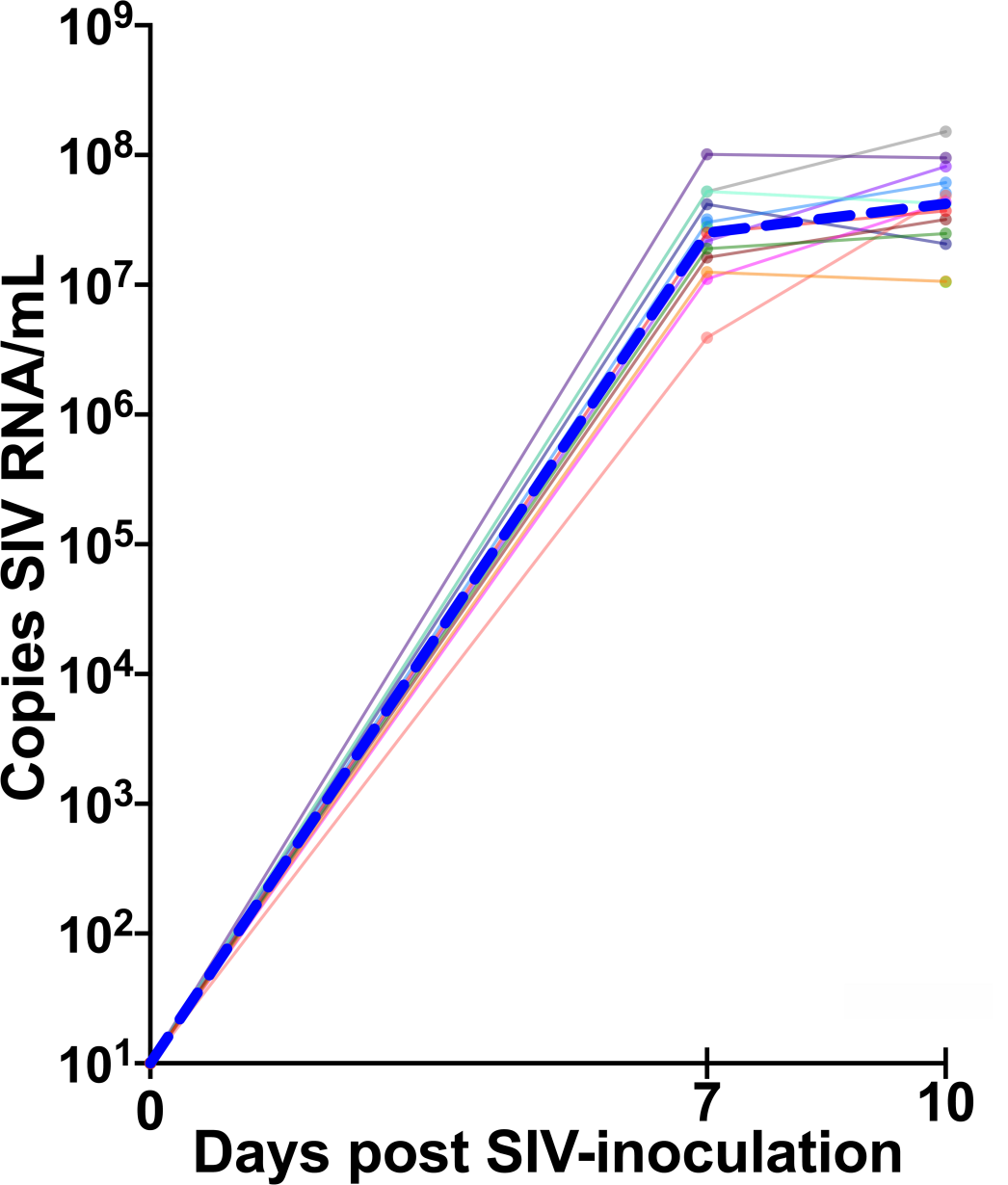
All 12 SIV-infected macaques had comparable high plasma viral loads during the acute phase of SIV infection with the peak group median plasma viral load (blue line) of 4.2 x10^7^ copies/mL at day 10 post-inoculation. Individual animal plasma viral loads are represented by the thin colored lines.

**Fig 6.**
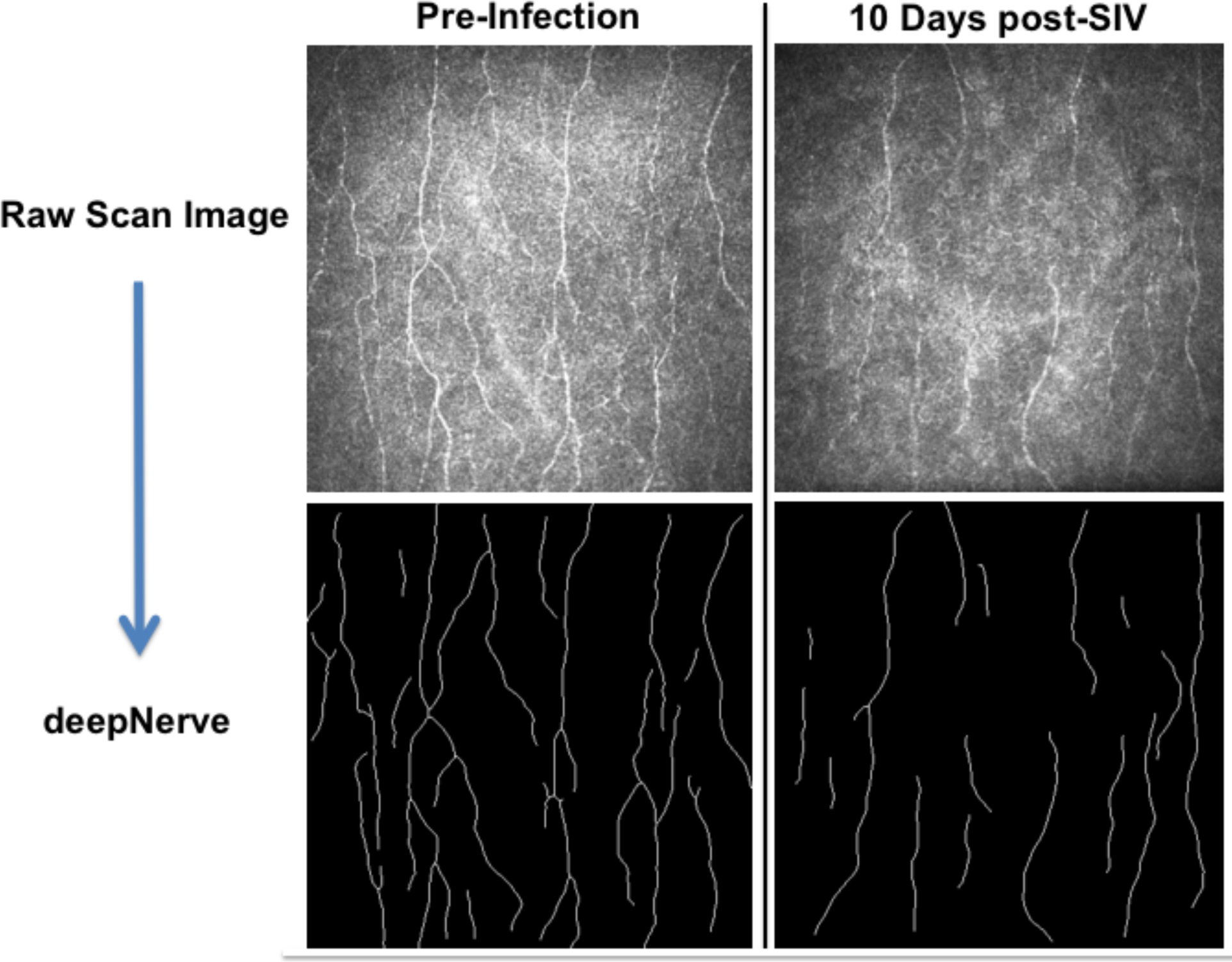
IVCM images (top) and deepNerve representations of the same images of corneal nerve fibers (below) from a pigtailed macaque at pre-infection (left column) and 10 days post-SIV infection (right column) illustrate loss of subbasal nerve fibers with SIV.

deepNerve-based measurements of subbasal nerve fiber features demonstrated that CNFL declined significantly by day 10 of infection (P= 0.01, paired t-test, Figure 7) with 10 of the 12 macaques having lower measurements at day 10 consistent with sensory nerve fiber loss. In contrast with the SIV-infected macaques, corneal nerve fiber length in uninfected control animals imaged at comparable repeat timepoints were not significantly different (P = 0.34; paired t-test). These data demonstrate the ability of the combination of IVCM and automated analysis to detect corneal nerve fiber loss during acute infection, an early timepoint when SIV replicates in sensory ganglia macrophages and induces both immune activation and metabolic alterations.(4, 7, 24)

**Figure 7.**
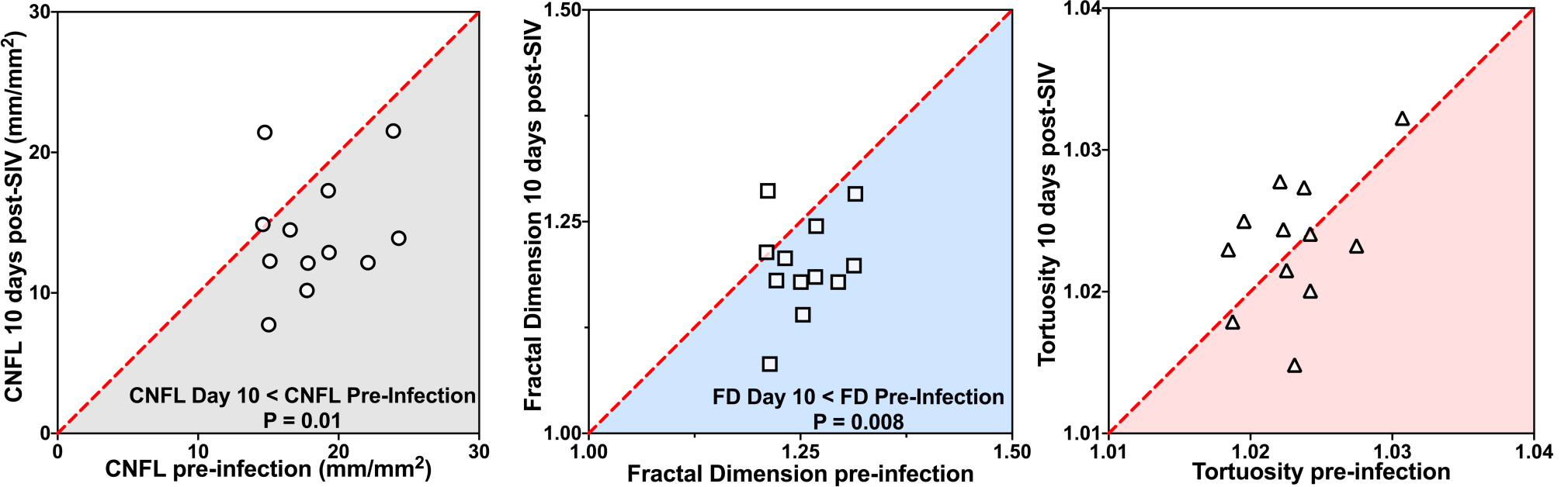
10 of the 12 SIV-infected macaques had lower corneal nerve fiber counts 10 days post-inoculation with SIV than their baseline pre-infection corneal scan values (left panel; P = 0.01, paired t-test). Similarly, fractal dimension declined with SIV infection in of 12 animals (middle panel; P = 0.008, paired t-test). In contrast, tortuosity was not altered during acute infection (right panel).

Similar to corneal nerve fiber length, fractal dimension also declined with SIV infection with 10 of the 12 animals with lower counts after SIV infection (Figure 7; P = 0.008, paired t-test). In contrast, corneal nerve fiber tortuosity was not significantly altered during acute infection and this may not be a sensitive marker of relatively acute damage.

## Discussion

Asian macaques are frequently studied to understand a variety of diseases including those that involve loss of small sensory nerve fibers associated with development of peripheral neuropathy.(19) Studies of both pigtailed macaques (*Macaca nemestrina*) and rhesus macaques (*Macaca mulatta*), for example, have been highly informative for understanding HIV pathogenesis including peripheral nerve damage that underlies development of HIV sensory neuropathy.(4-7, 24-26) The current clinical standard for measuring sensory nerve fiber loss in human and macaque subjects is performing counts of epidermal nerve fibers present in skin biopsies.(27) This technique is invasive and requires extensive tissue processing and analysis making repeat sampling for longitudinal assessments challenging. In contrast, non-invasive in vivo corneal confocal microscopy can quickly collect multiple images of the small sensory nerve fibers comprising the corneal subbasal plexus in a non-invasive manner, facilitating longitudinal acquisition of data. These images then can be rapidly evaluated using advanced automated processing approaches that objectively measure alterations in morphologic features.(28-33)

We have previously demonstrated corneal sensory nerve fiber loss in the SIV/pigtailed macaque model of HIV-induced PNS damage by studying corneal innervation *ex vivo* using immunohistochemical staining for the nerve marker βIII-tubulin followed by automated image analysis.(14) Corneal nerve fiber loss was well correlated with decline in epidermal nerve fibers. This study built on those findings by using IVCM to collect images of the macaque subbasal plexus, adapting techniques established in human studies.(12)

Using our well-characterized SIV/pigtailed macaque model of HIV-induced PNS damage, to determine whether acute SIV infection altered sensory nerves in the subbasal plexus, we obtained repeat IVCM images of 12 animals both pre-infection and then at 10 days post-infection time-points allowing us to directly compare corneal nerve fiber alterations in individual subjects. Automated analysis of IVCM images revealed that 10 of the 12 animals developed significant corneal nerve fiber alterations consistent with sensory nerve fiber damage and loss. Our earlier studies have shown that SIV replicates in macrophages in the trigeminal ganglia as early as 7 days post-infection.(4) Furthermore, we have used targeted transcriptomic analyses to reveal that immune activation and metabolic alterations can be detected in dorsal root ganglia 7 days post-SIV infection.(24) These changes in sensory ganglia during acute SIV infection likely underlie the sensory nerve fiber loss detected by both IVCM and skin biopsy-based measurements of epidermal nerve fiber density. Understanding acute alterations induced by HIV by studies in the SIV/macaque model is important because effective neuroprotective interventions are most valuable at the earliest stages of nerve damage.

A recent cross-sectional study showed that IVCM can detect corneal nerve fiber loss in people living with HIV.(13) A separate study of HIV cohorts demonstrated signs of neuropathy in 35% of individuals during the first months primary HIV infection before starting ART, suggesting that HIV causes early damage to the sensory nervous system as reported in SIV.(3) Future studies tracking corneal nerve fiber alterations in people living with HIV before and while on ART will contribute to our understanding of the pathogenesis of HIV neuropathy and enhance efforts to prevent and treat this debilitating disease.

To address the more challenging imaging conditions resulting from the aforementioned difference in macaque anatomy and lack of compliance with respect to human subjects, we developed a customized deep learning-based approach to assess macaque corneal nerve parameters in support of macaque IVCM studies. (16) We used this combination of IVCM and automated analysis to generate normative reference data sets for female and male pigtailed and rhesus macaques of varying ages. We did not detect significant species, sex, or age differences in measured corneal nerve parameters, however the approximately two-fold range in variation found in normal animals illustrates the challenge of performing cross-sectional studies if group sizes of animals are relatively small.

The ability to track corneal nerve fiber alterations longitudinally in individual animals by IVCM will enhance the rigor and reproducibility of macaque-based studies that are typically limited in size given animal availability constraints and high cost of macaque studies. Future studies will explore the potential of our deep learning-based approach for analyzing corneal nerve fiber properties in human subjects including those living with HIV to help guide prevention and clinical management of neuropathy.

## Acknowledgements

We thank Dr. Robert J. Adams, the JHU Retrovirus Lab team, and JHU Research Animal Resource members for their generous assistance in completing this project.

